# Targeting *Listeria* virulence-regulon glutathion-cavity by evolutecas, docking and Boltz2

**DOI:** 10.1101/2025.06.09.658548

**Authors:** J Coll

## Abstract

Computational explorations are described by screening and generating peptide and acetic-polymer evolutecas targeting one of the virulence regulons of *Listeria monocitogenes* (*Lm*). *Lm* are among the most important multiple-resistant food-borne infective bacteria whose antibiotic resistances raise worldwide concerns. The *Lm* glutathion cavity called **<u>p</u>**ositive **r**egulatory **f**actor **<u>A</u>** (prfA) virulence regulon, was computationally targeted here because of the glutathion-dependent prfA activation of many *Lm* virulent genes. Computationally docking screening of both tri-,tetra-, and penta-mer peptides or innovative acetic-polymers (mimicking amino acids) and evolutecas generating thousands of ligand candidates, were employed to optimize their fitting to glutathion-prfA cavities. Combining library-screenings and evoluteca generations, novel top-peptides and top-acetic-polymers were generated predicting both low-toxicities (to reduce any known undesirable side-effects) and low nanoMolar consensus affinities (to maximize their specificities). Co-evolved top-acetic-polymer smiles were also co-folded to prfA amino acid sequence (ligand-induced-fit) by deep-learning Boltz2. Comparative ADV and Boltz2 results confirmed similar targeted cavity and most top-ligand affinities, despite their sharply different algorithms. Whether some of the top affinities / conformations correspond to the most active prfA inhibitors will only depend on experimental validation. Therefore, some additional chemical synthesis and *in vitro* experimental validation, are strictly required to continue with the *Lm* prfA virulence explorations.

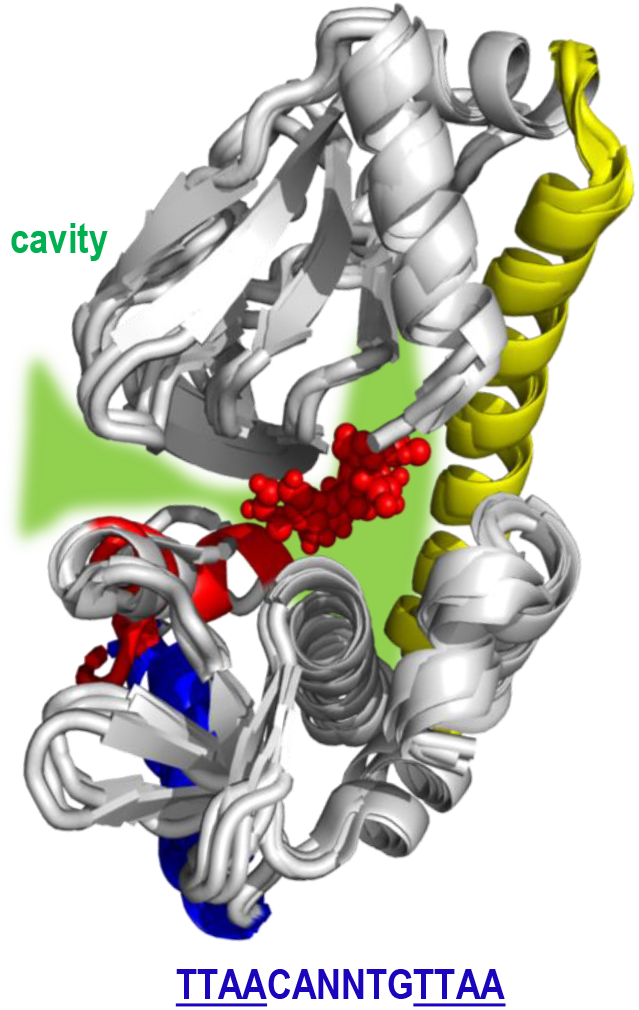

## The prfA regulon of *Lm* virulence

*Listeria monocitogenes* (*Lm*) are intracellular Gram-positive bacteria causing increasing worldwide concerns of these food-borne infections. The increasing emergence of hypervirulent *Lm* strains with pan-antibiotic resistances further limits any previously antibiotic-treatment options. Unexplored targets may be one of strategies to generate any new *Lm* virulent antibiotics.

Among computationally possible yet unexplored *Lm* targets, virulence regulons such as prfA (**p**ositive **r**egulatory **f**actor **<u>A</u>**) has been chosen here because it responds to the environment to induce the expression of hundreds of *Lm* virulence genes to generate biofilms, infection and antibiotic resistances.

The 237 amino acid prfA protein monomers becomes semi-activated by dimerization to weak binding to 14 bp palindromic DNA operators distributed among many virulent gene promoters (**Figure S3, blue letters: <u>TTAA</u>**CANNTG**<u>TTAA</u>**,). The tri-peptide glutathione (ECG in single letter amino acid code) allosterically stabilizes and activates the prfA-dimer DNA-binding.

Comparison of the crystallographic 3D structures of prfA in the ECG absence (2beo.pdb)^1^ and presence (5lrr.pdb)^2^, predicted its ECG-binding cavity **(Figure S3, green cavity)**. Each ECG tripeptide interacts with a prfA hydrophobic cavity formed by ∼ 60-70 (^63^Y, ^67^F), 110-125 (^126^Y) and other residues^2^.

Additionally, each prfA monomer contains an N-terminal domain (residues 1–108), a main long α-helix which interfaces with the other monomer (∼ 109–137), and one **<u>h</u>**elix-**t**urn-**<u>h</u>**elix DNA-binding HTH segment (170-196) similar to other prokaryotic DNA-binding proteins (flexible loop 170-177 + connecting loop 174-184 + α-helix 183-196)(**Figure S3, red cartoons**). The ECG or the prfA G^145^S mutation, induce α-helix conformations on the 170-177 prfA residues, stabilizing DNA-binding.

Although the binding affinity of ECG to prfA is ∼ 1000 nM, their eukaryote intracellular concentrations in host eukaryote cells are ∼ 10 mM, which could compensate such ECG low affinity. Nevertheless, the *Lm* glutathion synthase enzyme generates additional ECG tripeptide molecules if provided with any environmental Cystein and/or peptides-containing Cystein. The prfA 3D structures ± ECG are very similar having an overall carbon backbone rmsd difference of only 0.25-0.35 Å ^2^. However, in the presence of ECG, the prfA-DNA binding affinity increases to ∼ 50 nM. In contrast, to Cystein-containing peptides, no-Cysteine-containing peptides induce prfA inhibition through steric blockade of the ECG binding cavity^3^. The strength of the prfA DNA binding, depends on the ratio between Cystein-containing (activating) *versus* no-Cystein (inhibitory) peptides. Such mechanism may provide a control of expression of unneeded virulence when outside of a Cystein-containing environment. Thus, the ECG-mediated prfA conformational changes to activate DNA-binding can be inhibited by no-Cystein peptides such as LL (6hck.pdb) or EVFL (8cb7.pdb), which sterically blocked the ECG-binding cavity^3^. The hydrophobic side-chains of the ECG binding cavity suggest inhibitors of small molecular weight and highly-hydrophobicity. However, since particular peptides, could also lock the prfA in an activated conformation similar to the G^145^S mutation, experimental validation is critical to evaluate any possible inhibitions.

Therefore, the present computational search, was based on several working hypothesis including candidates which should:

i. contain no Cysteins.
ii. sterically block any ECG-binding cavities in an inactivating manner,
iii. have small molecular weight sizes to fit the ECG-binding cavity,
iv. predict higher docking affinities than ECG (i.e., low nanoMolar), and
v. require chemical synthesis for experimental validation

Compounds predicting the first four properties can be computationally predicted. However, the final requirement requires experimental validation.

To generate large numbers of candidates here, the strategy included:

a. screening tri-, tetra-, and penta-peptide possible libraries, without Cystein,
b. screening tri-acetic-polymers eliminating terminal Nitrogens and conserving amino acid side-chains,
c. generating evolutecas of thousands of ligands by the “build evolutionary library” co-evolutions of ***<u>D</u>****ata****<u>W</u>****arrior* ***<u>B</u>****uild* ***<u>E</u>****volutionary* ***<u>L</u>****ibrary* (DWBEL)^2-5^, and
d. Consensus affinity estimations by traditional docking with **<u>A</u>**uto**<u>D</u>**ock**<u>V</u>**ina (ADV) and deep-learning generation with most recent Boltz2.

DWBEL algorithms were employed before as one of the best alternative to generate thousands of ligands rather than screen extralarge drug-like banks^4, 5^ or predict earlier deep-learning docking models from protein sequences ^6-8^ or from recent RFdiffusion^9-13^. Based on our own preliminary RFdiffusion applications to snake venoms^14^ and monkeypox^15^, our attempts to generate RFdiffusion small peptides fitting the ECG cavity of *Lm* prfA were unsuccessful, most probably because their targeted cavities were inner at the prfA molecule and not in its surface (not shown). The DWBEL co-evolution alternative was/is a powerful method to generate large numbers of small molecular weight alternative candidates through mimicking natural evolution. In our hands, DWBEL had computationally generated higher affinity gains of non-toxic drug-like alternatives when targeting different protein-ligand pairs. For instance, new antibiotics for resistant *Staphilococcus*^16^, Abaucin-derivatives against resistant *Acinetobacter*^17^, non-human anticoagulant rodenticides^18^, monkeypox Tecovirimat-resistant mutants^19^, inner-cavity SARS omicron^20^, inflammatory coronavirus ORF8 protein^21^, prokaryotic models of human K^+^ channels^22^, inner-cavity of VHSV rhabdovirus^23^, malaria circumsporozoite protein^19^, RSV resistant mutations^24^, anti-HIV-Vif A3G^25^ and A3A hypermutational human metastasis^26^.

Combining computational screening of peptide and acetic-polymer libraries with co-evolutionary generated evolutecas of thousands of ligands, were only the first steps to propose new candidates for prfA inhibition of virulence. Docking of DWBEL children by traditional **A**uto**D**ock**V**ina (ADV) was then performed to generate consensus affinity predictions. An extension of consensus were undertaken by applying Boltz deep-learning recent methods^27,28,29^. In particular Boltz2 is different from traditional docking like ADV because their dynamic and simultaneous generation of best-fitting both protein and ligand. The Boltz2 and similar strategies^29^ are like co-folding, induced-fit or simultaneous modeling. Boltz2, generate *de novo* 3D structures from amino acids sequences and ligand smiles without any conformational information. Boltz2 predicts the protein 3D structures by Alphafold in the presence of each ligand conformer and outputs the best-fitting protein/ligand complexes. In contrast, DWBEL and ADV (or any other classical docking algorithms), sample for the best-docking ligand conformers to a fixed 3D structure protein. Therefore, in contrast to the results obtained by traditional docking, the conformation of the protein complexed with the ligand might depend on each ligand (ligand induced-fit). As recently anticipated by others the combination of traditional docking and deep-learned methodologies “*could yield synergistic performance gains*” ^29^. Exploration of de *novo* generations was initiated here by providing Boltz2 with thousands of DWBEL co-evolved non-toxic fitted-children generated from one selected top-acetic-polymer parent. Although the results confirmed similarities of both prfA targeted binding/docking cavities and some of their top-affinity ranked-order predictions, their protein/ligand conformations and the ranges of affinities greatly differed. To further understand the possible significance of all these predictions, chemical synthesis and *in vitro* experimental validation of *Lm* virulence and/or biological functions are required.

## Computational prediction results

### Screening of peptide and peptide-derivatives

Comparisons among the crystallographic *Lm* prfA monomers of apo^2^, and complexed with ECG^1^, EVFL^3^ and LL^3^, predicted that the inhibitory EVFL and LL no-Cystein peptides sterically blocked the conformational changes on the ∼170-177 residues, otherwise occurring in the presence of ECG and/or Cystein peptides (**Figure S3, red cartoons)**. Therefore, to compete with ECG-binding, screening for higher affinities was performed among tri-peptide libraries containing all except Cysteins. One library of 6859 possible tri-peptides excluding Cysteins, was generated by home-designed Python scripts and ChemAxom 3D converter. The ADV docking to apo prfA resulted in most tri-peptides predicting higher affinities than ECG (**Figure 1A, blue dashed-line**). The highly hydrophobic top-tri-peptides FWW, FWF, FFW predicted the highest ∼ 1 nM affinities (lower docking-scores) (**Figure 1A, cyan circles**). These predictions confirmed that the hydrophobicity favoured the affinities of possible competitive inhibitors to the ECG binding-cavity, as previously proposed from crystallographic studies^2^.

**Figure 1.**
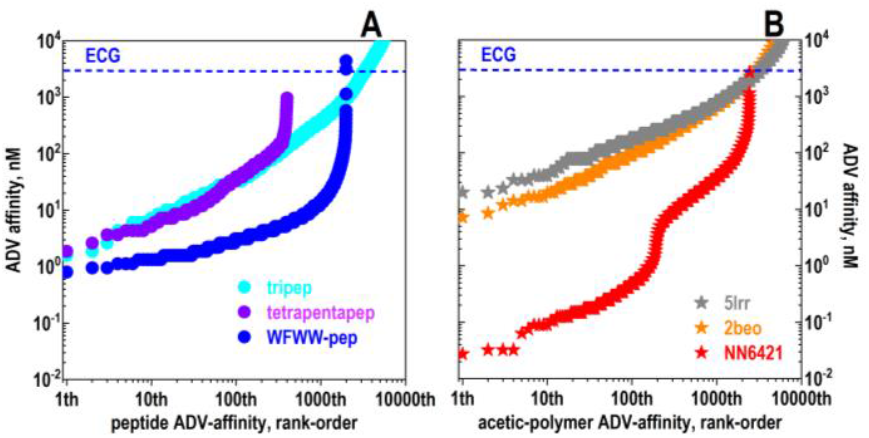
ADV-prfA docking rank profiles of non-Cystein peptides and tri-acetic derivatives. **A)** the 3-, 4- and 5-mer peptides were generated in single letter code by home-made Python scripts. The 4- and 5-mer computational predictions were mixed for simplification. The single amino acid letter code were converted to amino acid sdf files using a dictionary of amino acid smiles by the ChemAtom Converter web server (https://datascience.unm.edu/tomcat/biocomp/convert). **B)** Tri-acetic acid derivatives were generated by home-made Python scripts using a dictionary of single tri-acetic polymers corresponding to the amino acid smiles mostly by eliminating the amino of the 20 amino acids, excluding those corresponding to Cysteins. In both peptides and acetic-derivatives, their 2D conformations were maintained by generating their DW mmff94s+ conformers before ADV docking. Most of the resulting 3D conformers were ADV docked to 2beo.pdb (apo-prfA) monomers, except when indicated to 5lrr.pdb (ECG-conformation without ECG). PyRx ADV docked affinities were finally rank-ordered. **Blue dashed horizontal line**, ECG (glutathion) ADV binding affinity to 2beo.pdb **A)** 3-4-5-mer peptides (tripep, tetrapentapep) and DWBEL WFWW-derivatives to apo prfA 2beo.pdb **B)** 3-Acetic acid polymers and NN6421 DWBEL-derivatives to apo prfA 2beo.pdb and 5lrr.pdb±ECG **Cyan circles**, all possible 3-mer peptides, except Cystein docked to prfA (6859 peptides). **Violet circles**, all possible 4- and 5-mer peptides derived from FWW tripeptide (399 peptides) docked to prfA **Blue circles**, DWBEL derivatives from WFWW-tetra-peptide NN364 docked to prfA. **Gray stars**, all possible tri-acetic polymers docked to prfA 5lrr.pdb ± ECG (6859 polymers) **Orange stars**, all possible tri-acetic polymers docked to apo prfA 2beo.pdb (6959 polymers) **Red stars**, DWBEL derivatives from the top NN6421 tri-acetic polymer docked to apo prfA

All possible tetra-peptides derived from the FWW top-tri-peptide were then generated adding all possible N- and/or C-terminal amino acids, except Cysteins. The corresponding ADV computational predictions could not identify any significant affinity improvements of the top-tetra-peptides (**Figure 1A, violet-cyan circles**). However, PyMol visualization of their docked complexes revealed that most of the top-tetra-peptides coded for the same amino-terminal W (Tryptophan) and targeted the downward sites of the ECG-cavity. In contrast, the tri-peptides at their carboxy-terminal amino acids were variable (i.e., W,E,Y,G,D,A,I,Q) and targeted the upper sites of the ECG-cavity (not shown). These observations suggest that carboxy-terminal amino acids could be added to the tetra-peptides to improve their docking affinities. However, the addition of all possible amino acids, except Cystein to the C-terminal residues of WFWW, failed to generate significant affinity improvements (**Figure 1A, violet circles**). Representative top-penta-peptides were: WFWWW, WFWWE, WFWWY, WFWWG, WFWWD, WFWWA, WFWWI, WFWWQ.

### Evolutecas derived from tetra-peptides

To explore for additional tetra-peptide-derivatives predicting affinity improvements, their C terminus was extended by small atom combinations. For that the WFFW top-tetra-peptide structure parent was conserved during DWBEL partial co-evolutions performed of its carboxy-terminus carbon atom to derive WFFW-X children. For the fitting cavity selection during co-evolution, the prfA WFFW-E cavity (containing one of the largest cavities targeted by penta-peptides), was selected. Such partial co-evolutions, randomly generated 19023 raw-children of which 2011 were selected by fitting and filtered as non-toxic fitted-children WFFW-X derivatives. However, none of these co-evolved derivatives predicted significative affinity improvements when compared to the tri-, tetra- or penta-peptides (**Figure 1A, blue circles**). Despite conserving a high number of hydrophobic side-chains and multiple molecular variations at their C-terminus, no improvement of affinities were detected.

### Screening of acetic-polymers

To explore additional alternatives to computationally improve docking affinities to the ECG-binding cavity, innovations were developed by generating “tri-acetic polymers” with amino acid side-chains. The so called here tri-acetic polymers were similarly designed by constructing a dictionary of “no amino” acid-likes deleting terminal Nitrogens but containing the same 19 amino acid side-chains (all except Cystein). Their targeting to the cavities of the prfA monomers, predicted similar affinities whether the cavities were derived from the ±ECG models or the apo prfA (**Figure 1B, gray** and **orange stars**, respectively), but did not improve the ∼ 1-10 nM affinities obtained by the corresponding tri-peptides (**Figure 1A, violet-cyan circles**). However, PyMol visualization of top-tri-acetic polymers in complex with prfA, identified additional nearby amino acids targeting distal sites at the ECG-binding cavity (**Figure S1B**).

### Evolutecas derived from acetic-polymers

The results of the screening of acetic-polymers suggested that DWBEL co-evolution could help to generate derivatives predicting higher activities. Therefore, the top-NN6421 tri-acetic polymer (“WSW”-build with the corresponding amino acid side-chains) was selected as both parent molecule and prfA cavity to generate their co-evolution children. The DWBEL program randomly generated 41622 raw-children of which 2437 were selected as being non-toxic and cavity-fitted-children. The rank-ordered ADV affinites of the generated children predicted top non-toxic fitted-children increasing 100-1000-fold their affinities (**Figure 1B, red stars**). Several top-children predicted maximal affinities ∼ 0.03 nM (**Figure 1B red stars**), >100-fold higher than those obtained by the tri-, tetra-, penta-peptides and/or their evolved peptide-derivatives (**Figure 1A, cyan, violet, blue**). Because these novel acetic-polymer derivatives did not contained any Cystein side-chains and occupied longer prfA cavities (**Figure 2C,D**) than ECG (**Figure 2A**), these compounds may be sterically blocking ECG-binding. Possible *in vitro* side-effects may also be reduced, because such high affinities may allow their use at low concentrations. Nevertheless, compared with the peptides, tri-acetic derivatives may be more difficult to synthesize. Chemical synthesis and strict *in vitro* experimental validation remains to be done.

**Figure 2.**
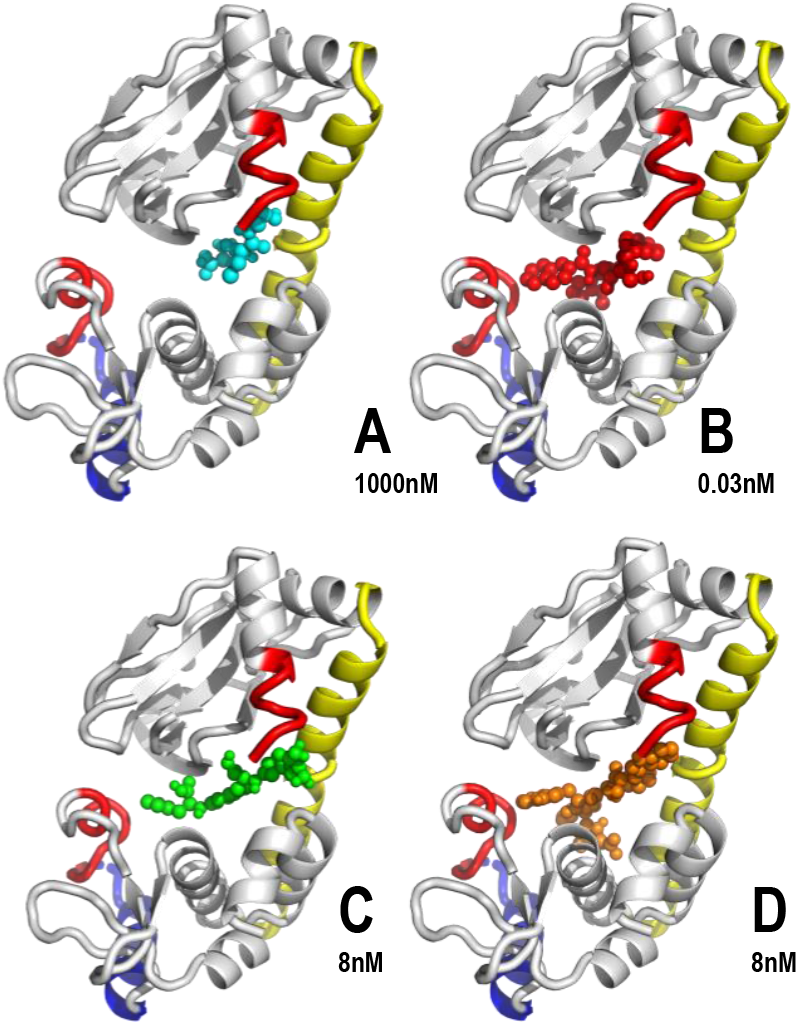
Visualization of docking cavities targeted by ECG (A), NN331 DWBEL-derivative (B),FWW (C) and WFWWE (D) to the apo prfA conformation. ADV affinities were approximated for C and D by their mean ± 1 nM (n=2), those for A and B were ± 500 nM and 0.01 nM (n=2), respectively. **A)** ECG. main cavity binding/docking ECG **B)** NN331 DWBEL-derived top-children from NN6421. **C)** tri-peptide FWW (NN1424). **D)** penta-peptide WFWWE (NN 364)

Top-peptides and top-acetic-polymers predicted higher affinities to prfA than the ECG affinity (∼ 500-fold or ∼ 30000 higher affinities, respectively) (**Figure 1AB, blue horizontal dashed line)**. The comparison of nearby amino acids of some representative top-peptides and top-acetic-polymer derivative conformers, predicted that all of them always included the two prfA main sites of the ECG-binding cavity (∼ 60-70 and 110-125 residues) (**Figure 3, ECG vertical cyan rectangles**). Additional prfA residues were targeted by both top-peptides or top-acetic-polymers such as the amino-terminal ∼10 first amino acids, 140-165 or amino acids around the 230 carboxy-terminal residue (**Figure 3, different colors**).

**Figure 3.**
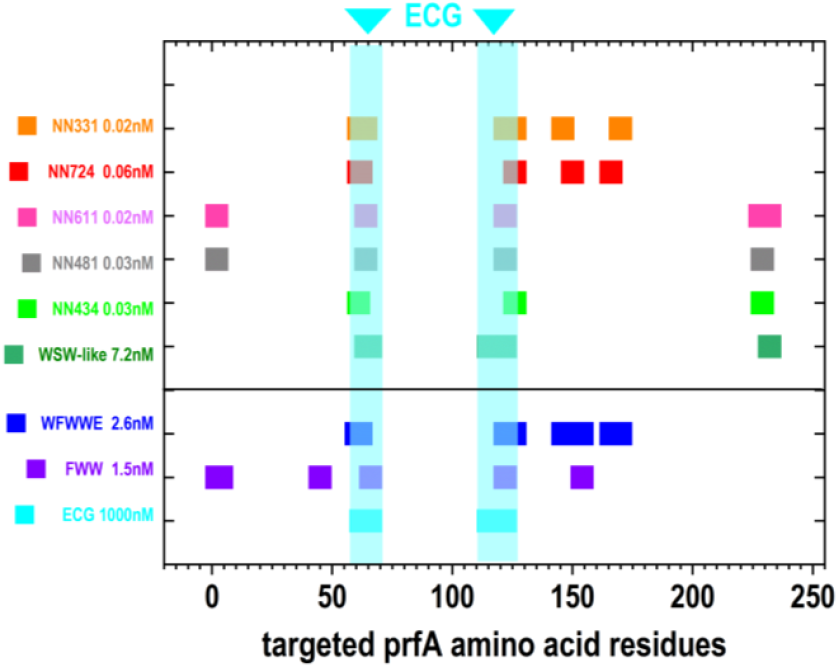
prfA Amino acids targeted by ECG and top-peptides and top-acetic-polymers. Nearby amino acid positions within 4 Å of docked top-molecules were computationally predicted by home-designed PyMol/Python *.py scripts and drawn in Origin templates. **X-axis**, amino acid position numbers of the prfA amino acid monomer (left amino to right carboxy positions). **Lower part of the Figure**, ECG and representative top-tri-(FWW) and penta-(WFWWE) peptides. **Upper part of the Figure**, WSW-like tri-acetic parent and selected representatives of its top-DWBEL co-evolved children ranked-ordered by their NN numbers. Their approximated affinities were calculated by ADV docking. **Cyan vertical rectangles**, prfA amino acids targeted by ECG docking. **Violet rectangles**, prfA amino acids targeted by docking of the tri-peptide FWW. **Blue rectangles**, prfA amino acids targeted by docking of the penta-peptide W**FWW**E (derived from FWW). **Other color rectangles**, prfA amino acids targeted by top-acetic-polymer derivatives (DWBEL from WSW-like tri-acetic-polymer)

These top-molecules are only some of the many other possible solutions that were identified among those predicting higher prfA affinities than ECG, suggesting they could successfully compete for binding to prfA. However, although easier to synthesize, top-peptides contain a large proportion of highly hydrophobic side-chains that may be difficult to handle and/or cause excesive unspecific bindings to other proteins. Given the existence of a vast chemical space with many other alternatives, perhaps the acetic-polymers predicting lower hydrophobicities with higher affinities, may better survive any experimental steps. Only *in vitro* experimental validation could demonstrate any possible *Lm* virulence inhibition.

### Deep-learning Boltz2 of NN6421 acetic-polymer derivatives

The ADV affinity predictions of the evoluteca generated from the top-NN6421 acetic-polymer were compared to the affinity predictions of Boltz2. The NN6421-derived DWBEL evoluteca consisted in an sdf file output containing 2437 non-toxic fitted-children (**Figure 1B, red stars**). Although alternative evolutecas could be generated from other parent molecules, the NN6421-derived DWBEL evoluteca biased model was selected because of its highest ADV affinities.

Two inputs were required to run Boltz2:

i. The prfA monomer amino acid sequence (single-letter code), and
ii. The sdf file output of 2437 non-toxic fitted-children.

Although Boltz2 often performs better when provided with 3D templates of the same or similar proteins (pdb or cif files), no template was provided here for prfA. This option was chosen because during *de novo* generation, Boltz2 always redesigns 3D structures, refine chains, build missing loops, remodel protein regions, etc. Furthermore, preliminary Boltz2 explorations using the 2beo.cif monomer as template, did not generate any affinity improvements (not shown). Therefore, no prfA templates were employed in the Boltz2 generations described here.

One of the Boltz2 ipynb versions most suitable to read and processes the DWBEL sdf file outputs was the ipynb modification described by Aarshit Mittal (https://colab.research.google.com/github/whis9/boltz2/blob/main/boltz2.ipynb). Additional home-designed code was introduced into that ipynb Colab to process thousands of ligands in batches, including reading the input sdf file, and their conversion to smiles before running Boltz.

Preliminary Boltz2 modellings were performed with top-children NN611, NN434, NN481, NN724 (**Figure 1, red stars** and **Figure S2**). Both ADV and Boltz2 predicted similar prfA targeted docking and “binding”-cavities for all top-children (**Figure 4, AB**). However, while ADV predicted highly similar top-children conformers, Boltz2 predicted different conformers for each of them.

**Figure 4.**
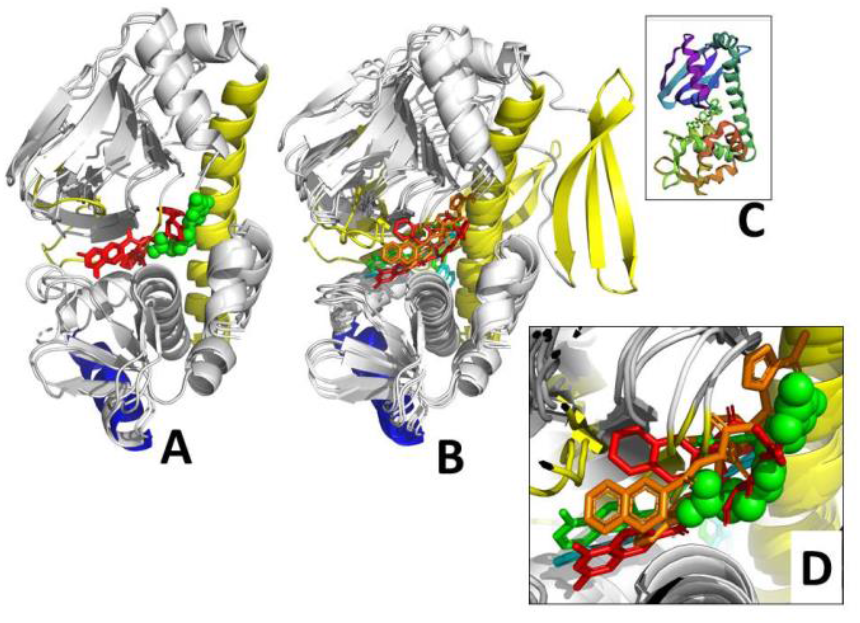
Comparison of DWBEL acetic-polymer top-conformers by ADV and Boltz2. Merged models corresponded to DWBEL generated top-children NN611, NN434, NN481, NN724 (**Figure S2**). The Boltz2.ipynb modified by A-Mittal further fine-tuned and home-modified was employed. Numbering of the PyMol prfA cartoons corresponded to the Boltz2 amino acid positions resulting from prfA-ECG. To note that all top-children and prfA conformers were very similar when docked by ADV. In contrast the Boltz2 changed all top-children conformers and NN434 and NN481 top-children induced important conformational changes on prfA ß-sheets (81-110 residues). **A**, crystallographic fixed prfA model + merged with DWBEL-ADV top-conformers predicting ∼ 0.03 nM affinities **B**, Boltz2 prfA modeled + merged Boltz2 top-conformers predicting 10-100 nM affinities **C**, Boltz modeled prfA-NN434 as it appeared in the Boltz2 ipynb **D**, Details of the Boltz modeled prfA-top-conformers from B **Gray cartoons**, prfA carbon backbones. **Blue cartoons**, prfA DNA-binding α-helices **Yellow cartoons**, prfA main α-helix (156-183) and ligand-induced ß-sheet conformations (81-110) **Green spheres**, Boltz2 generated prfA-ECG. **Cyan sticks**, NN611 (∼NN331) top-children conformer **Orange sticks**, NN434 top-children conformer. **Green sticks**, NN481 top-children conformer **Red sticks**, NN724 top-children conformer

Additionally, NN434 and NN481 predicted important conformational changes on prfA, specially those at its ß-sheet region (81-110 residues) (**Figure 4 B, yellow ß-sheets**). The predicted top-children affinities were also different, since while ADV predicted ∼ 0.03 nM high affinities (**Figure 3**), Boltz2 predicted most of their affinities between 10-100 nM and ∼ 1000 nM for NN611 (**Figure 5**).

**Figure 5.**
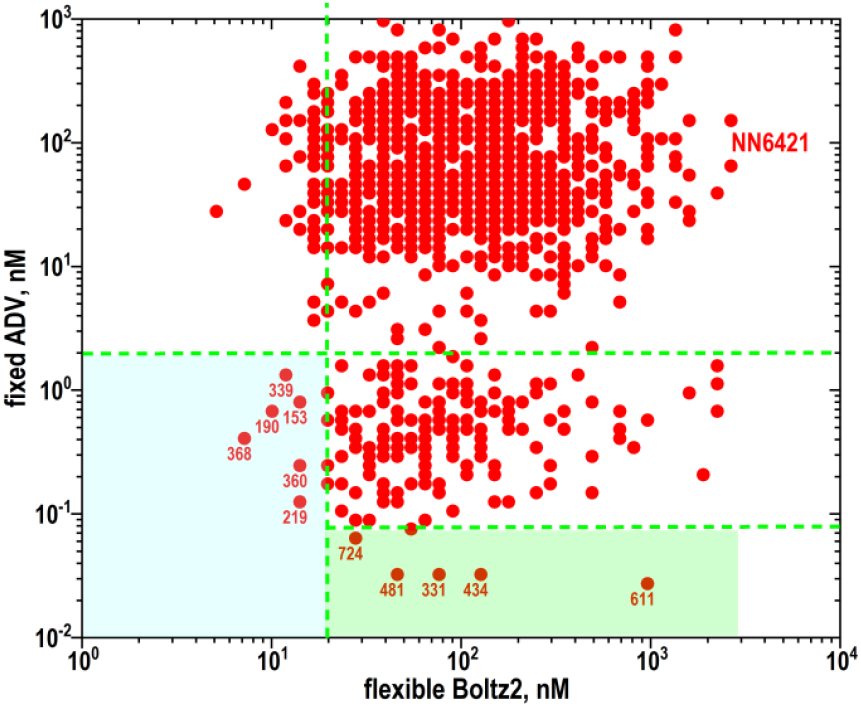
Comparison of predicted affinities between ADV and Boltz2. **DWBEL** randomly generated 41622 raw-children from the tri-acetic-polymer **NN6421** (upper right). These raw-children generated one evoluteca of 2437 children best-fitting the NN6421-prfA fixed-cavity (**Figure 1, red stars**). ADV and Boltz affinities were then calculated and the first DWBEL ∼ <NN1300 best-fitting children represented in this Figure. **ADV** docking searched for the best conformers of the DWBEL fitted-children to the prfA-ECG crystallographic fixed-cavity. Their outputs were reported in kcal/mol energy and converted to ∼ nM. **Boltz2** simultaneously generated *de novo* the best children conformers to the best prfA cavity conformation. Their outputs were reported in kcal/mol energy and converted to ∼ nM. The scales were arbitrary adjusted to improve the visual comparisons. **Green dashed lines**, arbitrary-thresholds for maximal affinities for ADV and Boltz **Red circles**, ADV *vs* Boltz2 predicted affinities of ∼3000 DWBEL fitted-children **Blue background** and **red numbers**, NN identification of new ADV and Boltz2 top-ligands. **Green background** and **red numbers**, NN724, NN481, NN331, NN434, NN611 were previously identified (**Figure 3**)

To further explore such affinity differences, 2473 non-toxic fitted-children from DWBEL-derived NN6421 (**Figure 1, red stars** and **Supplementary files /DWBELvsADVvsBoltz.zip**) were co-folded with prfA models by Boltz2. The **DWBEL** evolutecas were generated by selecting the non-toxic best-fitting children to one fixed-cavity within prfA. Their outputs were in negative docking scores that could not be converted to affinities. However most of the 1300 tops predicted very similarly low docking scores which were within an small compact DWBEL range (between ∼ -110 to -155). As expected by their fittings, those low ranges would correspond to the highest affinities.

**ADV** docking searched for the best conformers among the DWBEL non-toxic fitted-children by targeting similar prfA fixed-cavity than the one employed for DWBEL (ADV outputs were in kcal/mol energy and were converted to ∼ nM). In contrast, **Boltz2** *de novo* generated the DWBEL-children conformers best-fitting to the best prfA cavity conformation (Boltz2 outputs were in kcal/mol IC^50^ and were converted to ∼ nM). Despite their highly different algorithm strategies (**Table S3**), the best-fitted ligands targeted similar prfA cavities by both ADV and Boltz2 (**Figure 4 AB**). However, there were no apparent correlations between their affinities in the top-ranges studied (**Figure 5)**. Correlation trends were apparent between ADV and Boltz2 affinities corresponding to children > NN1300 predicting lower affinities than those represented (not shown).

Nevertheless, two groups of ADV and Boltz2 top-ligands could be identified. The first group predicted ∼ 0.02-0.1 nM highest ADV affinities (low docking-scores). Most of them were distributed at Boltz affinities from 20-100 nM, while NN611 predicted ∼ 1000 nM (**Figure 5, green rectangle backgrounds**). This group included all the top-children identified before, such as NN481, NN331, NN434, NN611 and NN724 (**Figure 3** and **Figure S2**), confirming most of those ADV top-children were also found among some of the Boltz2 higher affinities. Most of the second group predicted 0.1-1 nM ADV affinities (higher-docking scores) and ∼ 10 nM Boltz affinities (NN368, NN190, NN339, NN360, NN219, NN153) (**Figure 5, blue rectangle backgrounds**). Additionally, the ADV and Boltz low affinities of the NN6421 parent (**Figure 5, upper right**) confirmed that the DWBEL co-evolution generated numerous new children ligand structures some of which may predict increasing affinities by both ADV and Boltz (**Supplementary files /DWBELvsADVvsBoltz.zip**).

Among many other possibilities, the discrepancies of affinity estimations between ADV and Boltz may be due to different: scoring functions, molecular physics, trained data, and/or fixed *vs* flexible protein models (**Table S2**).

In contrast to the DWBEL-ADV predictions targeting crystallographic apo prfA fixed models, Boltz2 dynamically generated *de novo* prfA models which can be different from the crystallographic structure. Because for maximal DNA-binding activity prfA undergoes ECG-induced conformational changes, the flexible Boltz predictions might be closer to reality. However, it is also possible that neither ADV, nor Boltz2 could reflect accurately the real prfA-ligand interaction. Perhaps the prfA cavity may have more probabilities to be real because of the approximated coincidence of crystallography, ADV and Boltz2. On the other hand, the true prfA and ligand conformations and their resulting affinities and therefore their ranking order may be somewhere in between, best predicted by one of the models or even outside the identified ranges. On the other hand, perhaps the ligands predicting the maximal affinities by both methods (three counting DWBEL) could have more probabilities for biological activity (**Figure 5, green rectangle background**), but it remains to be proven. Without experimental data, it is not possible to identify which of the computational methods is the best predictor of real binding.

## Computational limitations

The DWBEL-ADV predictions are strictly limited to computational docking made by including numerous simplifying constrains (i.e., fixed 3D protein structure and cavity targets, absence of water interactions which may interfere the docking processes, exclusively rely on approximated maximal affinites using only one or two consensus docking programs, targeting one of the possible alternative cavities, etc). The Boltz predictions are learned from previous protein and ligand conformational data without any other constrains. However, both computational predictions require experimental validation of their possible biological functions. Before experimental validation some of the top-predicted ligand candidates need to be chemically synthesized. Among all the molecules predicted, small peptides may be the easiest to synthesize for *in vitro* experimental tests. Despite all those limitations, the predicted prfA inhibitor candidates have not been described before and may also provide examples for further explorations and development of similar or related computational strategies. Any drug-like inhibitors of *Lm* virulence remain strictly dependent on additional *in vitro* and *in vivo* experimental validation.

## Conclusions

New drug-like small molecular weight compounds predicting no-toxicities together with low-nanoMolar affinities to the ECG-binding cavity of *Lm* prfA monomeric models have been computationally generated. Novel ligand generation by DWBEL co-evolution and ADV docking confirmation were combined to identify novel peptides and acetic-polymer candidates. DWBEL and ADV employed targeted rigid crystallographic 3D models of prfA while sampling different ligand conformers for their best-fitting defined prfA cavities. Two different affinity approximations were employed to rank-order ligand conformers. A sharp different approach analysed possible interactions using only prfA amino acid sequence and DWBEL generated smiles computational representations as inputs for recent deep-learning Boltz2 algorithms. Boltz2 simultaneously predicted the best prfA 3D structures together with the best ligand conformer. Although traditional docking and deep-learning methodologies identified similar prfA cavities, they differed in their prfA and ligand conformations/conformers. It is not yet known which of the identified top candidates, if any, would best predict real binding. In the absence of any experimental data, it may be impossible to know now.

## Supporting information

L21_7.ipynb

DWBELvsADVvsBoltz.dwar

## Supplementary information

**Table S1.**
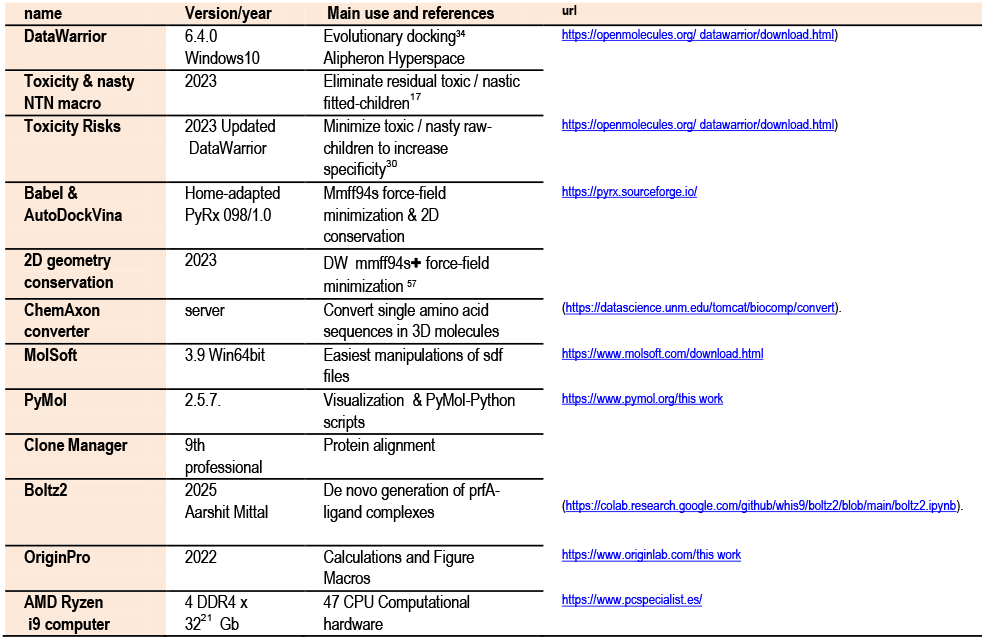
Computational software and hardware.

**Figure S1.**
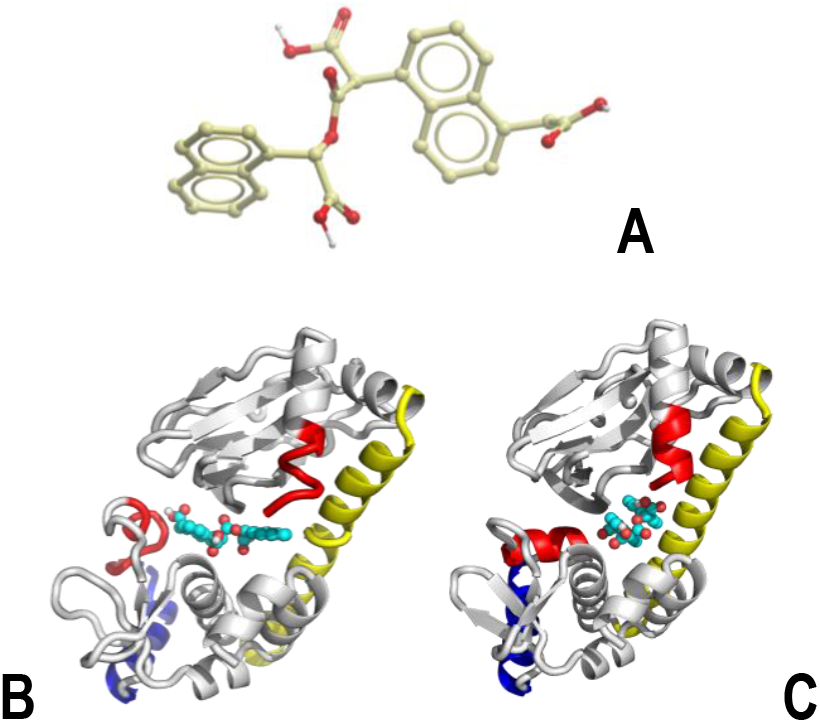
Top-acetic-polymer NN6421 (A) docked to apo prfA (B) and prfA±ECG (C) The top-tri-acetic-polymer NN6421 was docked to prfA monomers in the apo (2beo.pdb) and the ±ECG (5lrr without ECG). Comparison of NN6421 complexed to prfA in the apo (**B**) with the ±ECG (**C**) conformations could explain the sterically inhibition of NN6421 binding by the ECG induced 170-177 α-helix (**C, red α-helix**). In contrast, the NN6421 conformer was compact when docking to prfA in their ECG-induced/empty cavities (Figure S1C) **Grey cartoons**, carbons of the prfA monomer. **Yellow cartoons**, main α-helices 108-137 residues. **Blue cartoons**, partial DNA-binding HTH 183-196 residues. **Red cartoons**, prfA 170-177 residues changing after ECG binding. **Different color spheres**, derivative NN6421.

**Figure S2.**
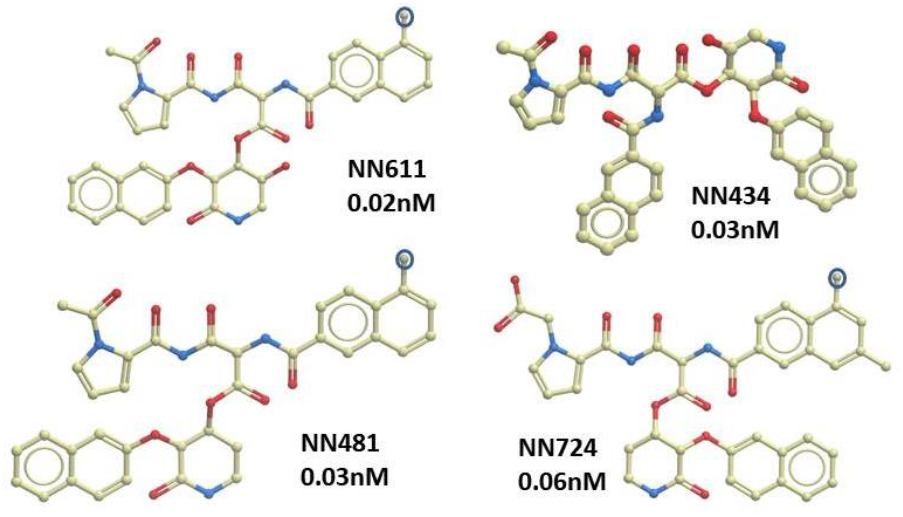
NN6421-derivative top-children. Examples of the similar 2D structures of some of the most representative top-NN6421 DWBEL derivatives which may deserve chemical synthesis for strictly required experimental validation steps. **Red atoms**, Oxygen. **Blue atoms**, Nitrogen, **Greenish atoms**, carbon. **Encircled greenish atoms**, Fluor.

**Figure S3.**
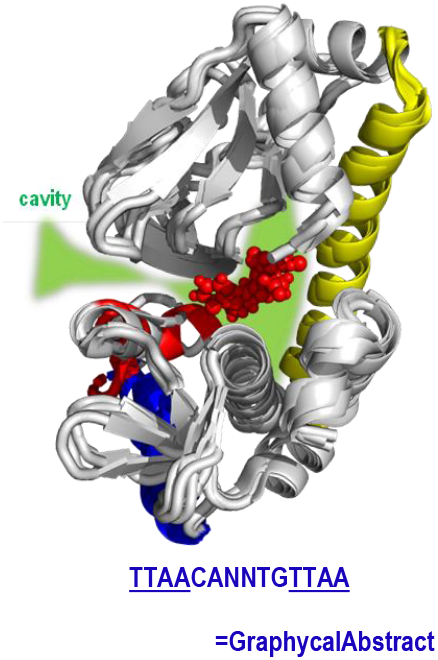
PyMol merged *Lm* prfA monomers: 2beo (apo-prfA), 5lrr (prfA+ECG), 8cb7 (prfA+EVFL) and 6hck (prfA+LL). **Grey cartoons**, 2beo.pdb. 5lrr.pdb, 8cb7, 6hck carbon backbones **Yellow cartoons**, 108-137 main α-helices. **Blue cartoons**, 183-196 DNA-binding HTH α-helices. **Green background**, possible alternative docking cavities around the crystallographic ECG-binding cavity (upper site). **Red spheres**, merged EVFL, LL inhibitory peptides and ECG. To note the steric contacts with the ECG-induced red α-helix. **Red cartoons**, 5lrr ECG-induced α-helix (∼170-177 residues), that includes stabilization of DNA-binding affinity. in contrast, the corresponding sequences in the presence of inhibitory peptides and in the apo-prfA showed variable conformations. **Blue letters**, operator DNA of 14-bp palindromic consensus sequence of *Lm* virulence genes.

**Table S3.**
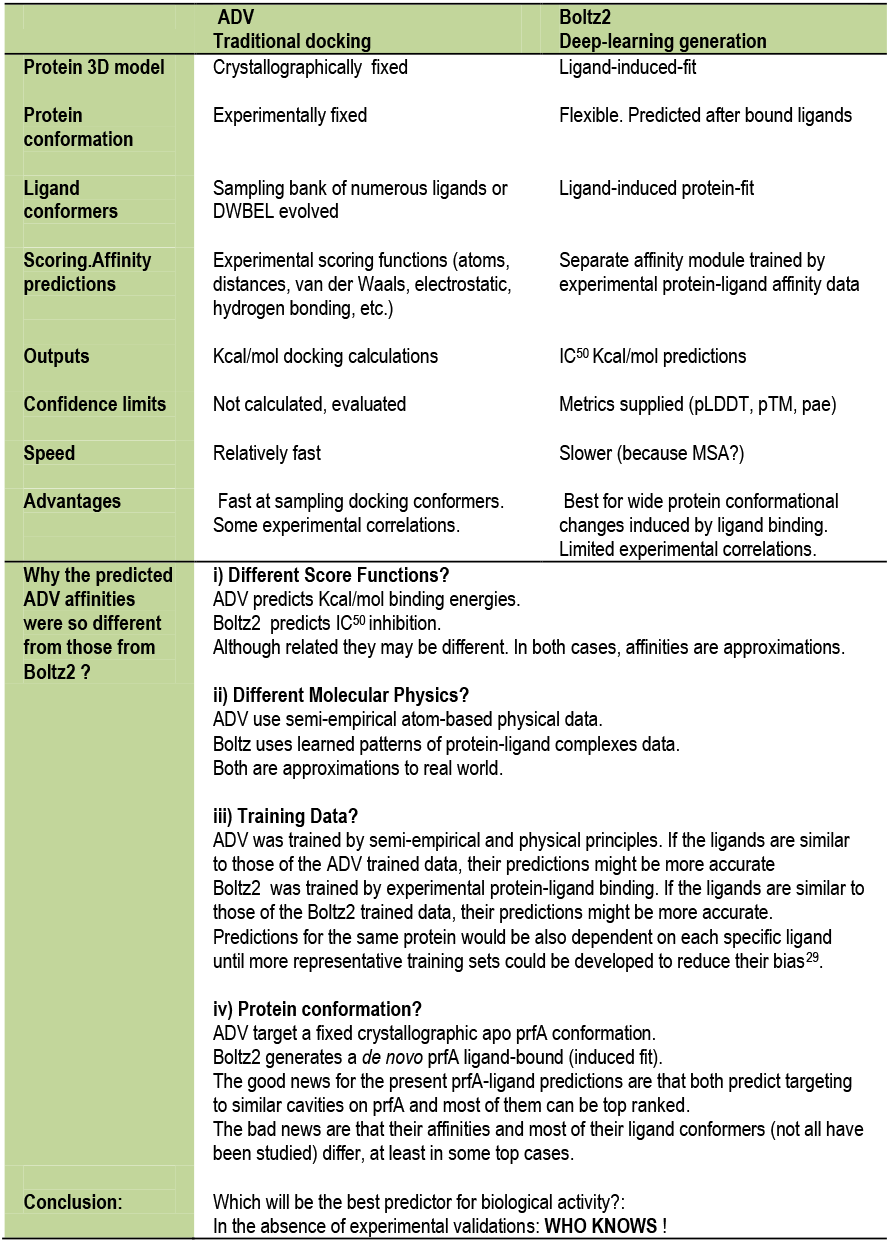
Comparison between ADV and Boltz2.

|  | ADV<br>Traditional docking | Boltz2<br>Deep-learning generation |
| --- | --- | --- |
| **Protein 3D model** | Crystallographically fixed | Ligand-induced-fit |
| **Protein conformation** | Experimentally fixed | Flexible. Predicted after bound ligands |
| **Ligand conformers** | Sampling bank of numerous ligands or DWBEL evolved | Ligand-induced protein-fit |
| **Scoring/Affinity predictions** | Experimental scoring functions (atoms, distances, van der Waals, electrostatic, hydrogen bonding, etc.) | Separate affinity module trained by experimental protein-ligand affinity data |
| **Outputs** | Kcal/mol docking calculations | IC<sub>50</sub> Kcal/mol predictions |
| **Confidence limits** | Not calculated, evaluated | Metrics supplied (pLDDT, pTM, pae) |
| **Speed** | Relatively fast | Slower (because MSA?) |
| **Advantages** | Fast at sampling docking conformers. Some experimental correlations. | Best for wide protein conformational changes induced by ligand binding. Limited experimental correlations. |
| **Why the predicted ADV affinities were so different from those from Boltz2 ?** | **i) Different Score Functions?**<br>ADV predicts Kcal/mol binding energies. Boltz2 predicts IC<sub>50</sub> inhibition. Although related they may be different. In both cases, affinities are approximations.<br>**ii) Different Molecular Physics?**<br>ADV use semi-empirical atom-based physical data. Boltz2 uses learned patterns of protein-ligand complexes data. Both are approximations to real world.<br>**iii) Training Data?**<br>ADV was trained by semi-empirical and physical principles. If the ligands are similar to those of the ADV trained data, their predictions might be more accurate. Boltz2 was trained by experimental protein-ligand binding. If the ligands are similar to those of the Boltz2 trained data, their predictions might be more accurate. Predictions for the same protein would be also dependent on each specific ligand until more representative training sets could be developed to reduce their bias<sup>29</sup>.<br>**iv) Protein conformation?**<br>ADV target a fixed crystallographic apo prfA conformation. Boltz2 generates a *de novo* prfA ligand-bound (induced fit). The good news for the present prfA-ligand predictions are that both predict targeting to similar cavities on prfA and most of them can be top ranked. The bad news are that their affinities and most of their ligand conformers (not all have been studied) differ, at least in some top cases. |  |
| **Conclusion:** | Which will be the best predictor for biological activity?: In the absence of experimental validations: **WHO KNOWS !** |  |

## Supplementary files

**- Lm21_7.ipynb**. The ipynb version modified by Aarshit Mittal (https://colab.research.google.com/github/whis9/boltz2/blob/main/boltz2.ipynb), was selected to run Boltz2. Additional home-designed code to batch affinity predictions, are still preliminary and in development. The actual version included:

**0)** Similar installation of dependencies, verify installation, import libraries, core Boltz interface and visualization functions. Additional it was provided by a first cell to mount the local Drive for automatic saves. Different version errors appeared, but did not interfere with the rest of the code. In our ColabPro environment this step have to be repeated to bypass the errors stop,

**1)** To define user-variables for Boltz2 running (job_name, batch_selection_input, batch size, drive_folder_path, protein_sequence, input sdf file) and to convert 2437 children sdf-file to their ligand NN name and smiles,

**2)** To generate batch size, the ligands were divided in a number of batches according to the total number of ligands and the user-desired batch_size (100 ligands were preferred),

**3)** Run the number of the batch_selection_input (in our ColabPro environment, running two simultaneous independent batches of 100 ligands each, took ∼ 9 hours to complete), and

**4)** After finishing the run, automatically export to Drive all batch_affinity_ results to csv, to avoid colab inactivity crashes and to allow easy export of the results to the local computer and graphic representation by Origin.

- **DWBELvsADVvsBoltz.dwar**. DW table containing 1473 children derived from NN6421. It was provided with threshold slider-filters to their DW docking-scores, ADV affinities and Boltz2 afinities. The DW table included molecular weights and clogP properties of the putative candidates. By moving up-down the slider-filters at the table’s right, the best-fitting children to particular threshold combinations could be selected. The dwar fil e can be opened by downloading DataWarrior free access at https://openmolecules.org/datawarrior/download.htm

## Computational methods

### PrfA 3D models

The amino acid sequences of *Lm* prfA were extracted from 5lrr.pdb (in complex with the ECG glutathion). Other prfA structure files were the apo-prfA (2beo.pdb), prfA+ECG cofactor (5lrr.pdb), prfA+EVFL inhibitory non-Cystein tetrapeptide (8cb7.pdb) and prfA+LL inhibitory non-Cystein dipeptide (6hck.pdb).

### Generation of peptide and acetic-polymer libraries

One text library of all possible 6859 tri-peptides, was generated by a home-made Python script from an smiles 20 natural amino acid dictionary combining all possible orders of three amino acids, except Cystein^3^. The initial single letter code tri-peptide text file was converted to one sdf file coding for atom tri-peptide structures by the Converter ChemAxon server (https://datascience.unm.edu/tomcat/biocomp/convert). To conserve their 2D geometries during ADV docking, 3D-conformers were generated for each tri-peptide by the DW mmff94s**+** force-field. All possible tetramers conserving the top FWW tri-peptide were generated adding all possible N- and/or C-terminal amino acids except C and following similar procedures than for the tri-peptides. Additionally, all possible pentamers were also generated from the top WFWW tetramer. Tri-acetic polymers were computationally generated by derivation of the single amino acid smiles dictionary except Cysteins, mainly by eliminating the amino-terminal peptide bond (tri-acetic derivatives) using appropriated home-made Python scripts.

### AutoDockVina docking

Before docking, any libraries in sdf files were converted to 3D-conformers applying the DW mmff94s**+** force-field algorithm^31^ to strictly preserve their 2D geometries during minimization, pdbqt generation and ADV docking.

Computational affinity predictions and prfA targeted cavities were performed by **<u>A</u>**uto**<u>D</u>**ock**<u>V</u>**ina (ADV)^46^ docking using the OpenBabel mmff94s^17^ force-field algorithm included into the home-tuned PyRx-0.98/1.0 packages. ADV was chosen because of their limited but relatively high accuracies compared to other docking programs experiencing world-wide ongoing improvements^10,29^. In particular ADV docking was employed here to:

1. Select the best parents and prfA targeted cavities by blind-docking (targeting grids of 30×30×30 Å, surrounding most of the prfA molecule),
2. Quantify the approximated affinities of the best-conformers in ∼ nM. The ADV predicts score outputs estimated in Kcal/mol^32,33-35^. Kcal/mol estimations were converted to ∼ nM affinities by applying the formula, 10^9^*(exp^(−Kcal/mol/0.592)^), and
3. identify nearby amino acids to ADV docked 3D conformer-prfA complexes to confirm their PyMol visual targeted prfA cavities.

### Co-evolutionary random-generation and selection of fitted-children molecules

Co-evolutionary docking was performed by **<u>D</u>**ata**<u>W</u>**arrior-**<u>B</u>**uild **<u>E</u>**volutionary **<u>L</u>**ibrary (DWBEL). DWBEL co-evolutions randomly generated molecular atom variations during 3 consecutive-independent runs. For each run, selection of children used fixed criteria values and relative weights (x1-x4) such as minimal docking-score (4 value) and maximal relative importance weight (x4), optimal molecular weight <= 600 g/mol (x2), low-medium hydrophobicity LogP <=4 (x1), and minimal Toxicity risk <=1 (x4). From the parent molecule, DWBEL co-evolutions randomly added/inserted small atom variations to originate tens of thousands of consecutively ID numbered raw-children. The raw-children selected for best cavity-fitting and absence of know toxicity motifs, were finally filtered by a the NTN DW macro to exclude hundreds of children molecules containing any remaining signatures of mutagenesis, tumorigenicity, reproductive interference, irritant molecular signatures, and/or nasty motifs^19,36^. The resulting non-toxic fitted children were provided with NN numbers ordered from their highest to lower docking-scores, their mmff94s+ conformers generated and all the generated data rows (one children per row) saved to an unique sdf vs3 file. The final children were ADV docked and rank-reordered by their predicted ADV nM affinities.

### Evaluation of thousands of ligands by deep-learning Boltz2

The 2438 acetic-polymer children generated from NN6421 DWBEL co-evolution were docked by ADV (**Figure 2B, red star**) and their affinity predictions compared to those from Boltz2^27,28,29^. The ipynb version modified by Aarshit Mittal was selected for Boltz2. (https://colab.research.google.com/github/whis9/boltz2/blob/main/boltz2.ipynb)

Additional home-designed code to batch affinity predictions, are still on preliminary development (**Supplementary files/L21_7.zip**) including fine-tuning of:

**0)** Similar installation of dependencies, verify installation, import libraries, core Boltz interface and visualization functions, but including a first cell to mount the local Drive to provide automatic saves (different errors appear, but do not interfere with the rest of the code),

**1)** To define user-variables for Boltz2 running (job_name, batch_selection_input, batch size, drive_folder_path, and protein_sequence, input sdf file) and to convert the ligand sdf-files from DWBEL co-evolution to their name and smiles,

**2)** To generate batch size, the ligands are divided in a number of batches according to the total number of ligands and the user-desired batch_size,

**3)** Run the number of the batch_selection_input (in our environment of, ColabPro running 2 simultaneous batches of 100 ligands each, took ∼ 8 hours to complete the two independent batches), and

**4)** After finishing the run, automatically export to Drive all batch_affinity_ results to csv, to avoid colab inactivity crashes and to allow easy export of the results to local Origin.

Although this ipynb code was developed for comparison purposes, it could also be used to Boltz2 screening of many ligand smiles.

## Authors’ contributions

JC designed, performed and analyzed the work and drafted the manuscript. **Acknowledgements:**Thanks are due to Dr.Juan Arques at CSIC-INIA for suggesting the beginning of this work. Dr. Jo W for his valuable help with the acetic-polymers. Thanks are also due to the many people actively participating in the DW forum (https://openmolecules.org/forum/index.php?t=finduser&btn_submit=Find&), the Slack-Boltz forum (https://boltz-community.slack.com/ssb/redirect) and the Alphafold Discord forum https://discord.com/channels/. I must thank also to the Gemini /chatGTP machines for their valuable help with many coding challenges.

