## Supplementary material for "Targeting *Listeria* virulence-regulon glutathion-cavity by evolutecas, docking and Boltz2": L21_7.ipynb: - L21_7.ipynb

```
# %% [markdown]
[link text](https://)### *BOLTZ L21 Google Colab Interface*
2025-06-13 by Aarshit Mittal  
This notebook provides a simple way to harness the power of [BOLTZ 2](https://github.com/jwohlwend/boltz) (MIT license) for:
Protein+ligand-batch binding affinity calculatios
2025-06-21 modified by Julio Coll: focused on affinity estimation of ligand batches

# %% [markdown]
#**INSTALLATIONS:**

# %% [code]
#@title mount drive to colab
from google.colab import drive
drive.mount('/content/drive')
print("Google Drive mounted !")

# %% [code]
#@title Install Dependencies
print("to avoid possible errors by unrequired dependencies, run twice the Install Dependencis")
import os
import sys

# --- 1. Define required versions ---
# This makes it easy to update versions in the future
REQUIRED_TORCH_VERSION = "2.4.0"
REQUIRED_PL_VERSION = "2.5.0"

# A flag to determine if we need to run the installation
needs_install = False

# --- 2. Check if the correct versions are already installed ---
try:
    import torch
    import pytorch_lightning as pl

    # Check if the installed versions match our requirements.
    # We use .startswith() to ignore suffixes like "+cu118"
    torch_ok = torch.__version__.startswith(REQUIRED_TORCH_VERSION)
    pl_ok = pl.__version__.startswith(REQUIRED_PL_VERSION)

    if torch_ok and pl_ok:
        print(f"✓ Correct versions already installed.")
        print(f"  - PyTorch: {torch.__version__}")
        print(f"  - PyTorch-Lightning: {pl.__version__}")
        print("Skipping installation.")
    else:
        print("Mismatched versions detected. Reinstallation is required.")
        if not torch_ok:
            print(f"  - Found PyTorch {torch.__version__}, but require {REQUIRED_TORCH_VERSION}")
        if not pl_ok:
            print(f"  - Found PyTorch-Lightning {pl.__version__}, but require {REQUIRED_PL_VERSION}")
        needs_install = True

except ImportError:
    # This block runs if one of the libraries isn't installed at all
    print("Required libraries not found. Installation is required.")
    needs_install = True

# --- 3. Run installation and restart ONLY if needed ---
if needs_install:
    print("\nUninstalling existing PyTorch to avoid conflicts...")
    # Using sys.executable ensures we use the correct pip
    !{sys.executable} -m pip uninstall torch torchvision torchaudio -y -q

    print("Installing specified library versions...")
    !{sys.executable} -m pip install torch=={REQUIRED_TORCH_VERSION} torchvision==0.19.0 --index-url https://download.pytorch.org/whl/cu118 -q
    !{sys.executable} -m pip install pytorch-lightning=={REQUIRED_PL_VERSION} boltz py3Dmol -q
    print("Installation complete.")

    # Crucial step: Restart the runtime to load the new libraries
    print("\nIMPORTANT: Runtime is restarting to load new versions. Please wait...")
    os.kill(os.getpid(), 9)

# %% [code]
#@title Verify Installation
import torch
import pytorch_lightning as pl
import boltz
import py3Dmol

print("✅ Dependencies loaded successfully!")
print(f"✓ PyTorch version: {torch.__version__}")
print(f"✓ PyTorch-Lightning version: {pl.__version__}")
print(f"✓ BOLTZ installed")
print(f"✓ py3Dmol installed")
print("-" * 30)
print(f"✓ GPU: {torch.cuda.get_device_name(0) if torch.cuda.is_available() else 'Not available'}")

# %% [code]
#@title Import Libraries
import os
import sys
import subprocess
import json
import tempfile
import yaml
import shutil
from pathlib import Path
import time
from datetime import datetime
from typing import Dict, List, Tuple, Optional

import numpy as np
import pandas as pd
import matplotlib.pyplot as plt
import seaborn as sns
import py3Dmol
from google.colab import files
from IPython.display import display, HTML
import torch

# Configure environment
os.makedirs('output', exist_ok=True)
plt.style.use('seaborn-v0_8-darkgrid')
sns.set_palette("husl")

# Check GPU availability
device = torch.device('cuda' if torch.cuda.is_available() else 'cpu')
gpu_info = torch.cuda.get_device_name(0) if torch.cuda.is_available() else 'CPU only'
print(f"Device: {device} ({gpu_info})")

# %% [code]
#@title Core BOLTZ Interface

class BOLTZPredictor:
    """Main interface for BOLTZ 2 predictions"""

    def __init__(self, output_dir: str = "output"):
        self.output_dir = Path(output_dir)
        self.output_dir.mkdir(exist_ok=True)

    def run_prediction(self, input_data: dict, job_name: str,
                      use_msa: bool = True, verbose: bool = False) -> dict:
        """
        Execute BOLTZ prediction

        Args:
            input_data: YAML-compatible input dictionary
            job_name: Unique identifier for this job
            use_msa: Whether to use MSA server
            verbose: Print detailed output

        Returns:
            Dictionary containing results and file paths
        """
        # Create temporary YAML file
        with tempfile.NamedTemporaryFile(mode='w', suffix='.yaml', delete=False) as f:
            yaml.dump(input_data, f)
            yaml_path = f.name

        # Build command
        output_path = self.output_dir / job_name
        cmd = ["boltz", "predict", yaml_path, "--out_dir", str(output_path)]

        if use_msa:
            cmd.append("--use_msa_server")

        try:
            # Execute prediction
            process = subprocess.Popen(cmd, stdout=subprocess.PIPE,
                                     stderr=subprocess.PIPE, text=True)
            stdout, stderr = process.communicate()

            if process.returncode != 0:
                return {"success": False, "error": stderr or stdout}

            # Parse results
            result_dirs = list(output_path.glob("boltz_results_*"))

            files = {"pdb": [], "cif": [], "json": []}
            for result_dir in result_dirs:
                files["pdb"].extend(list(result_dir.rglob("*.pdb")))
                files["cif"].extend(list(result_dir.rglob("*.cif")))
                files["json"].extend(list(result_dir.rglob("*.json")))

            # Load JSON results
            results = {}
            for json_file in files["json"]:
                try:
                    with open(json_file) as f:
                        results[json_file.name] = json.load(f)
                except Exception as e:
                    if verbose:
                        print(f"Warning: Could not read {json_file.name}: {e}")

            return {
                "success": True,
                "output_dir": str(output_path),
                "files": files,
                "results": results,
                "stdout": stdout if verbose else None
            }

        except Exception as e:
            return {"success": False, "error": str(e)}
        finally:
            Path(yaml_path).unlink(missing_ok=True)

    def extract_metrics(self, results: dict) -> dict:
        """Extract key metrics from BOLTZ results"""
        metrics = {}

        for filename, data in results.items():
            if "confidence" in filename and isinstance(data, dict):
                metrics.update({
                    "confidence_score": data.get("confidence_score", 0),
                    "ptm": data.get("ptm", 0),
                    "iptm": data.get("iptm", 0),
                    "plddt": data.get("complex_plddt", 0)
                })
            elif "affinity" in filename and isinstance(data, dict):
                # FIXED: Properly interpret affinity results
                # affinity_pred_value is log10(IC50 in μM) - this is what BOLTZ outputs
                log_ic50_uM = data.get("affinity_pred_value", 0)

                # Convert log10(IC50 in μM) to IC50 in μM
                ic50_uM = 10 ** log_ic50_uM

                # Convert to nM (1 μM = 1000 nM)
                ic50_nM = ic50_uM * 1000

                # Convert to M for proper pIC50 calculation
                ic50_M = ic50_uM * 1e-6

                # Calculate standard pIC50 (negative log of IC50 in molar units)
                # pIC50 = -log10(IC50 in M)
                pic50 = -np.log10(ic50_M) if ic50_M > 0 else 0

                # Calculate binding free energy in kcal/mol
                # Using the formula from BOLTZ documentation: (6 - affinity) * 1.364
                # Note: this assumes affinity_pred_value is in the range where 6 corresponds to 1 μM
                delta_g_kcal = (6 - log_ic50_uM) * 1.364

                # Approximate Kd from IC50 (Kd ≈ IC50/2 for competitive inhibitors)
                # This is a rough approximation!
                kd_uM = ic50_uM / 2
                kd_nM = kd_uM * 1000
                kd_M = kd_uM * 1e-6

                # Calculate pKd from Kd in molar units
                pkd = -np.log10(kd_M) if kd_M > 0 else 0

                metrics.update({
                    # Raw BOLTZ output
                    "boltz_affinity_value": log_ic50_uM,  # This is log10(IC50 in μM)

                    # IC50 values
                    "log_ic50_uM": log_ic50_uM,
                    "ic50_uM": ic50_uM,
                    "ic50_nM": ic50_nM,
                    "pic50": pic50,  # Standard pIC50

                    # Kd approximations
                    "kd_uM": kd_uM,
                    "kd_nM": kd_nM,
                    "pkd": pkd,

                    # Energy
                    "delta_g_kcal": delta_g_kcal,

                    # Binding probability
                    "affinity_prob": data.get("affinity_probability_binary", 0)
                })

        return metrics

# Initialize predictor
predictor = BOLTZPredictor()

# %% [code]
#@title Visualization Functions

def visualize_structure(structure_path: Path, width: int = 800, height: int = 600):
    """Display 3D molecular structure"""

    with open(structure_path, 'r') as f:
        content = f.read()

    file_format = 'pdb' if str(structure_path).endswith('.pdb') else 'cif'

    view = py3Dmol.view(width=width, height=height)
    view.addModel(content, file_format)
    view.setStyle({'cartoon': {'color': 'spectrum'}})
    view.setBackgroundColor('white')
    view.zoomTo()

    return view

def plot_confidence_distribution(plddt_file: Path) -> plt.Figure:
    """Plot per-residue confidence distribution"""

    data = np.load(plddt_file)
    plddt_values = data['plddt'] * 100  # Convert to percentage

    fig, ax = plt.subplots(figsize=(10, 6))

    # Histogram
    n, bins, patches = ax.hist(plddt_values, bins=50, alpha=0.7,
                               color='#2E86AB', edgecolor='black')

    # Color code by confidence level
    for i, patch in enumerate(patches):
        if bins[i] < 50:
            patch.set_facecolor('#D32F2F')  # Red - Very low
        elif bins[i] < 70:
            patch.set_facecolor('#F57C00')  # Orange - Low
        elif bins[i] < 90:
            patch.set_facecolor('#FBC02D')  # Yellow - Confident
        else:
            patch.set_facecolor('#388E3C')  # Green - High

    ax.axvline(plddt_values.mean(), color='red', linestyle='--', linewidth=2,
               label=f'Mean: {plddt_values.mean():.1f}%')

    ax.set_xlabel('pLDDT Score (%)', fontsize=12)
    ax.set_ylabel('Number of Residues', fontsize=12)
    ax.set_title('Per-Residue Confidence Distribution', fontsize=14, fontweight='bold')
    ax.legend()
    ax.grid(axis='y', alpha=0.3)

    plt.tight_layout()
    return fig

# %% [markdown]
## **:-)  ⏬ ===========================   😊**

# %% [markdown]
# **1 -----VARIABLES FOR RUNNING & INPUT SDF-FILE to  NAME+SMILES --------**

# %% [code]
#@title variables for running & input sdf-file to name+smiles
job_name = "Lm21" #@param {type:"string"}
batch_selection_input = 24  #@param {type:"raw"}
selected_batch_index = int(batch_selection_input)-1
batch_size = 100  #@param {type:"integer"} #number of ligands per batch
drive_folder_path = "\"/content/drive/My Drive/Lm21\" " #@param {type:"string"}
# Define the protein sequence (same for all ligands)
protein_sequence = "NAQAEEFKKYLETNGIKPKQFHKKELIFNQWDPQEYCIFLYDGITKLTSISENGTIMNLQYYKGAFVIMSGFIDTETSVGYYNLEVISEQATKLTSISENGTIMNLQYYKGAFVIMSGFIDTETSVGYYNLEVISEQATAYVIKINELKELLSKNLTHFFYVFQTLQKQVSYSLAKFNDFSINGKLGSICGQLLILTYVYGKETPDGIKITLDNLTMQELGYSSGIAHSSAVSRIISKLKQEKVIVYKNSCFYVQNLDYLKRYAPKLDEWFYLACPATWGKLN" #@param {type:"string"}
#print(protein_sequence)
display(protein_sequence)
# Install RDKit if you don't have it
try:
    from rdkit import Chem
except ImportError:
    print("RDKit not found. Installing...")
    !pip install rdkit -q
    from rdkit import Chem
import logging
from rdkit import RDLogger
    # Disable RDKit warnings that appear
lg = RDLogger.logger()
lg.setLevel(RDLogger.ERROR)
#------------------------------------------------------------------------------------------------------
from google.colab import files
print("Please upload your SDF file containing the ligands.")
uploaded = files.upload()
uploaded_filename = list(uploaded.keys())[0]
sdf_file_path = f"/content/{uploaded_filename}"
print(f"File '{uploaded_filename}' uploaded successfully.")
if uploaded:
  ligands_to_process = []
  uploaded_filename = list(uploaded.keys())[0]
  sdf_file_path = f"/content/{uploaded_filename}"
    # sanitize=False is used in case there are problematic molecules
    # removeHs=False is used to keep explicit hydrogens if they are important
  suppl = Chem.SDMolSupplier(sdf_file_path, sanitize=False, removeHs=False)
  print("\nProcessing molecules from SDF...")
  for i, mol in enumerate(suppl):
      if mol is None:
          print(f"Warning: Could not read molecule {i+1}. Skipping.")
          continue
      try:
          # Attempt to get a common name property
          ligand_id = mol.GetProp('_Name')
      except KeyError:
          # If '_Name' is not found, try to get all properties and look for something like 'ID'
          props = mol.GetPropsAsDict()
          ligand_id = props.get('ID', f'Ligand_{i+1}') # Default to a generated name if no ID property is found
          print(f"Warning: '_Name' property not found for molecule {i+1}. Using ID '{ligand_id}' (or generated name) instead.")
      except Exception as e:
           print(f"Warning: Could not retrieve name for molecule {i+1}: {e}. Using generated name.")
           ligand_id = f'Ligand_{i+1}'
      # Generate the canonical SMILES string
      try:
          smiles = Chem.MolToSmiles(mol, isomericSmiles=True)
          # Format the name as "NN" followed by the ligand_id
          formatted_name = f"NN{ligand_id}"
          ligands_to_process.append({"name": formatted_name, "smiles": smiles})
          print(f"  Added {formatted_name} with SMILES: {smiles}")
      except Exception as e:
          print(f"Warning: Could not generate SMILES for ligand '{ligand_id}' (molecule {i+1}): {e}. Skipping.")
  print(f"\nFinished processing sdf file. Created list with {len(ligands_to_process)} ligands.")
  print(ligands_to_process)


# %% [markdown]
# **2 -------- GENERATE BATCH SIZE --------**

# %% [code]
#@title Generate Batch-size
#batch_selection_input = 7  #@param {type:"raw"}
#selected_batch_index = int(batch_selection_input)-1
#batch_size = 100  #@param {type:"integer"} #number of ligands per batch
if 'ligands_to_process' in locals() and ligands_to_process:
    num_ligands = len(ligands_to_process)
    num_batches = (num_ligands + batch_size - 1) // batch_size
    ligand_batches = []
    for i in range(num_batches):
        start_index = i * batch_size
        end_index = min((i + 1) * batch_size, num_ligands)
        ligand_batches.append(ligands_to_process[start_index:end_index])
    print(f"Total ligands loaded: {num_ligands}")
    print(f"Generated 1-{num_batches} batches of ~ {batch_size} ligands each")
    print(f"BATCH_SELECTION_INPUT: 1 to {num_batches}")
    # Display batch information and prepare for selection
    batch_options = [(f"Batch {i+1} ({len(batch)} ligands)", i) for i, batch in enumerate(ligand_batches)]
elif 'ligands_to_process' not in locals():
    print("Error: 'ligands_to_process' list not found. Please run the SDF loading cell first.")
else:
     print("Warning: 'ligands_to_process' list is empty. No batches generated.")

# %% [markdown]
# **3 -------- RUN THE NUMBER OF THE BATCH_SELECTION_INPUT --------**

# %% [code]
#@title run the number of the batch_selection_input
#batch_selection_input = 7  #@param {type:"raw"}
#selected_batch_index = int(batch_selection_input)-1
if 'ligand_batches' in locals() and selected_batch_index >=0 and selected_batch_index < len(ligand_batches):
    current_batch = ligand_batches[selected_batch_index]
    print(f"RUNNING BATCH {selected_batch_index + 1} ({len(current_batch)} ligands)")
    all_ligand_results = []
    from IPython.display import display, Javascript # Import Javascript
    # Print header before the loop
    print(f"{'Ligand':<9} {'Kcal/mol':<9} {'IC50 nM':<9}")
    print("-" * 31) # Separator line
    for ligand_info in current_batch:
        ligand_name = ligand_info["name"]
        ligand_smiles = ligand_info["smiles"]
        current_job_name = f"batch_{selected_batch_index+1}_{ligand_name}" # Add batch number to job name
        input_data = {
            "version": 1,
            "sequences": [
                {
                    "protein": {
                        "id": "A", # Protein ID
                        "sequence": protein_sequence.upper().strip(),
                    }
                },
                {
                    "ligand": {
                        "id": "B", # Ligand ID
                        "smiles": ligand_smiles.strip()
                    }
                }
            ],
            "properties": [
                {
                    "affinity": {
                        "binder": "B" # Specify the ligand ID as the binder
                    }
                }
            ]
        }
        # Run prediction for the current ligand
        result = predictor.run_prediction(input_data, current_job_name, use_msa=True)
        if result["success"]:
            metrics = predictor.extract_metrics(result["results"])
            metrics['ligand_name'] = ligand_name
            metrics['batch'] = selected_batch_index + 1 # Add batch number
            all_ligand_results.append(metrics)
            if "boltz_affinity_value" in metrics:
              # Print values under the header
              print(f"{ligand_name:<9} {metrics.get('delta_g_kcal', 0):<9.1f} {metrics.get('ic50_nM', 0):<9.1f}")
            else:
                 print(f"{ligand_name:<9} {'N/A':<9} {'N/A':<9}") # NA Indicate no results
        else:
            print(f"Prediction failed for {ligand_name}: {result['error']}")
            display(Javascript("window.scrollTo(0, document.body.scrollHeight);")) #scroll down
        # Optional: Add a small delay between ligands to avoid overwhelming the server/resource
        # time.sleep(5) # Sleep for 5 seconds between ligands
    print(f"\n--- Batch {selected_batch_index + 1} Processing Complete ---")


# %% [markdown]
# **4 -------- EXPORT TO DRIVE ALL-BATCH AFFINITY-RESULTS TO CSV --------**

# %% [code]
#@title export to drive all-batch affinity-results to one csv file
import os
import shutil
from datetime import datetime
from google.colab import files
import pandas as pd
from pathlib import Path
drive_folder_path = "/content/drive/My Drive/Lm21" #@param {type:"string"}
def export_selected_batch_results(batch_index: int, results_df: pd.DataFrame, drive_path: str):
    if results_df.empty:
        print(f"No results found for batch {batch_index}. Nothing to export.")
        return
    # Ensure the target directory exists in Google Drive
    target_dir = Path(drive_path)
    target_dir.mkdir(parents=True, exist_ok=True)
    print(f"Saving results to: {target_dir}")
    desired_cols = ['ligand_name', 'delta_g_kcal', 'ic50_nM', 'kd_nM', 'affinity_prob', 'confidence_score']
    existing_desired_cols = [col for col in desired_cols if col in results_df.columns]
    if not existing_desired_cols:
        print(f"None of the requested columns found in the results for batch {batch_index}. Nothing to export.")
        return
    filtered_df = results_df[existing_desired_cols].copy() # Use .copy() to avoid SettingWithCopyWarning
    numeric_cols_to_format = [col for col in existing_desired_cols if col != 'ligand_name' and filtered_df[col].dtype in ['float64', 'float32', 'int64', 'int32']]
    format_dict = {col: '{:.1f}' for col in numeric_cols_to_format}
    for col, fmt in format_dict.items():
      filtered_df[col] = filtered_df[col].map(fmt.format)
    timestamp = datetime.now().strftime("%Y%m%d_%H%M%S")
    csv_filename = f'batch_{batch_index}_ligand_affinity_results_{timestamp}.csv'
    csv_filepath = target_dir / csv_filename # Save to the specified Drive folder
    try:
        filtered_df.to_csv(csv_filepath, index=False)
        print(f"Created CSV file: {csv_filename}")
        print(f"Size: {csv_filepath.stat().st_size / 1024:.1f} KB")
        # No need to call files.download() as we are saving to Drive
        print(f"Results for Batch {batch_index} saved to Google Drive.")
    except Exception as e:
        print(f"Error saving file to Google Drive: {e}")
# Make sure these variables are available from the previous execution
if 'selected_batch_index' in locals() and 'all_ligand_results' in locals():
    # Create a DataFrame from all_ligand_results
    results_df = pd.DataFrame(all_ligand_results)
    export_selected_batch_results(selected_batch_index + 1, results_df, drive_folder_path)
elif 'selected_batch_index' in locals() and 'current_batch' in locals():
    # If only current_batch is available, process results from the last executed batch
    results_df = pd.DataFrame(all_ligand_results) # Assuming all_ligand_results was populated during the last batch run
    export_selected_batch_results(selected_batch_index + 1, results_df, drive_folder_path)
else:
    print("Could not find necessary variables ('selected_batch_index', 'all_ligand_results', or 'current_batch'). Please run the batch processing cell first.")
```
